# Sustainable Production of Camptothecin from an *Alternaria* sp. isolated from *Nothapodytes nimmoniana*

**DOI:** 10.1101/2020.08.23.263129

**Authors:** I. A. H. Khwajah Mohinudeen, Rahul Kanumuri, K. N. Soujanya, R. Uma Shaanker, Suresh Kumar Rayala, Smita Srivastava

**Author notes:** K. N. Soujanya current affiliation: JSS college for women (Autonomous), Saraswathipuram, Mysuru-570009, India.

## Abstract

A total of 132 endophytic fungal strains were isolated from different plant parts (leaf, petiole, stem and bark) of *Nothapodytes nimmoniana*, out of which 94 were found to produce camptothecin in suspension culture. *Alternaria alstroemeriae* (NCIM1408) and *Alternaria burnsii* (NCIM1409) demonstrated camptothecin yields up to 426.7±33.6 μg/g DW and 403.3±41.6 μg/g DW, respectively, the highest reported production to date. Unlike the reported product yield attenuation in endophytes with subculture in axenic state, *Alternaria burnsii* NCIM1409 could retain and sustain the production of camptothecin up to ~200 μg/g even after 12 continuous subculture cycles. The camptothecin biosynthesis in *Alternaria burnsii* NCIM1409 was confirmed using ^13^C carbon labelling (and cytotoxicity analysis on different cancer cell lines) and this strain can now be used to develop a sustainable bioprocess for *in vitro* production of camptothecin as an alternative to plant extraction.

## INTRODUCTION

Camptothecin (1), first reported in 1966,^1^ is a monoterpene indole alkaloid. It is commercially produced from plants, mainly *Camptotheca acuminata* and *Nothapodytes nimmoniana.*^2^ Camptothecin is the third most in-demand alkaloid after taxol and vinca-alkaloids for anti-cancer applications, and inhibits DNA topoisomerase I in cancer cells leading to cell death.^3^ Among several camptothecin derivatives, topotecan, irinotecan and belotecan were approved and marketed as anti-cancer drugs, while others including silatecan, cositecan, lipotecan, simmitecan, chimmitecan, exatecan, namitecan, lurtotecan, elomtecan, diflomotecan, gimatecan, tenifatecan and genz-644282 are under various stages of clinical investigation. ^4,5^ Owing to the high demand of camptothecin, both the plants *C. acuminata* and *N. nimmoniana* are endangered and overharvested.^6^ In the Western Ghats of India, complete trees of *N. nimmoniana* are uprooted for camptothecin extraction. To produce 1 ton of camptothecin, around 1000-1500 tons of *N. nimmoniana* wood chips are needed.^7^ Hence, the annual demand for *N. nimmoniana* plants has been more than 1000 tons in recent years.^6^ Thus, to prevent such plant sources from extinction and to meet the ever-increasing demand of camptothecin, an alternative means of camptothecin production is needed.

It is well known that the endophytes can have the potential to produce their host plant-based metabolites. Since the discovery of the first taxol producing endophyte,^8^ endophytes in general have gained attention as a potent alternative source of plant metabolites. Though several successful attempts have been made to isolate endophytes, the commercial production of bioactive compounds using endophytes has not yet been established. The bottleneck has been the inconsistent product yield, which is generally reported to be lost with successive subculture under axenic state.^9^ Many camptothecin producing endophytes have been reported in the last decade. However, the yields reported have been low (0.5-45 μg/g) and inconsistent with subculture in axenic state (Table 1).

**Table 1.**
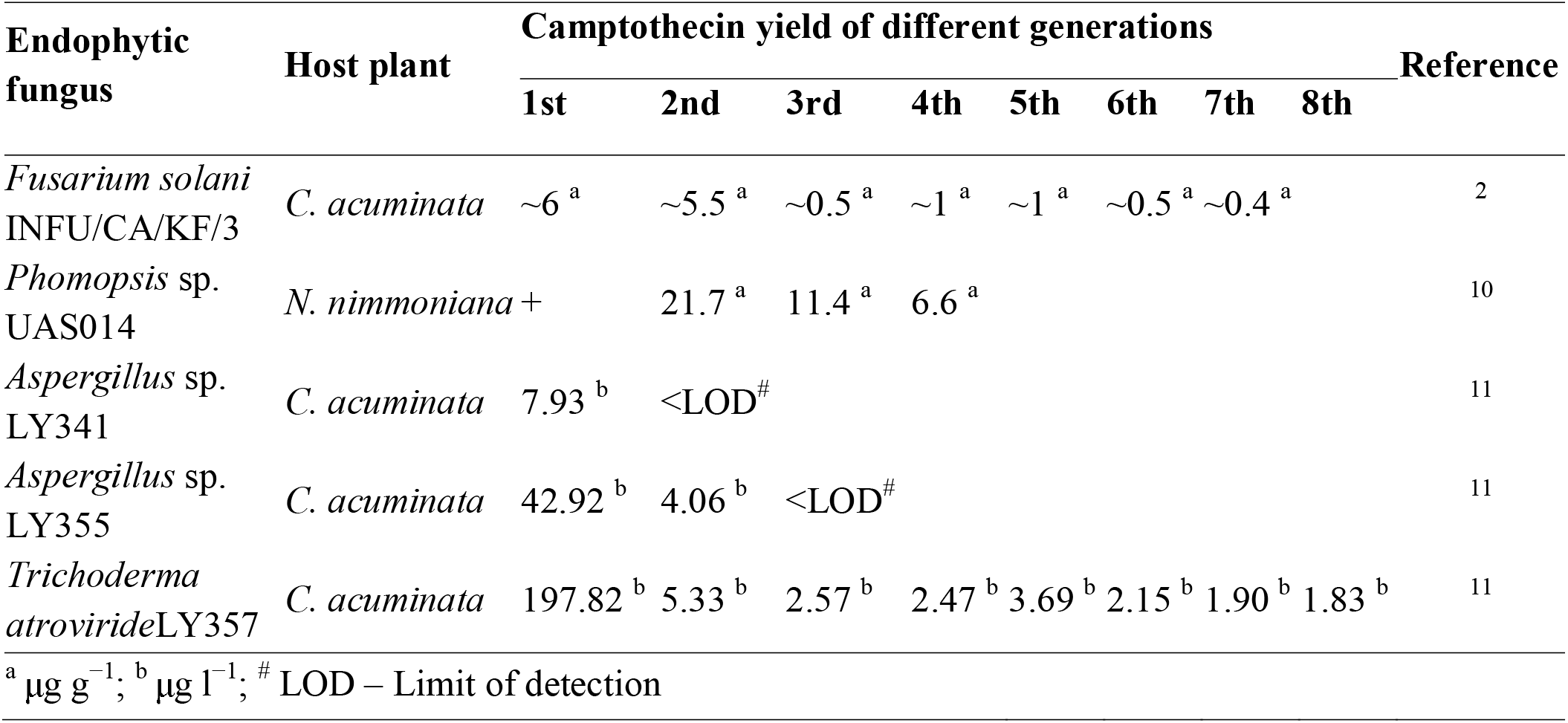
Reports on the attenuation of camptothecin produced from endophytes

In this study, we isolated a high camptothecin yielding endophyte from *N. nimmoniana,* demonstrating sustainable production of camptothecin in suspension culture. Increased relative abundance of ^13^C labelled camptothecin molecules in the axenic culture extracts, when the fungus was cultivated in the presence of D-[U-^13^C]-glucose, demonstrated inherent biosynthesis of camptothecin in the isolated organism. Also, the cytotoxic effect of the crude extract of camptothecin from the fungal suspension culture was demonstrated on colon, lung, ovarian and breast cancer cell lines.

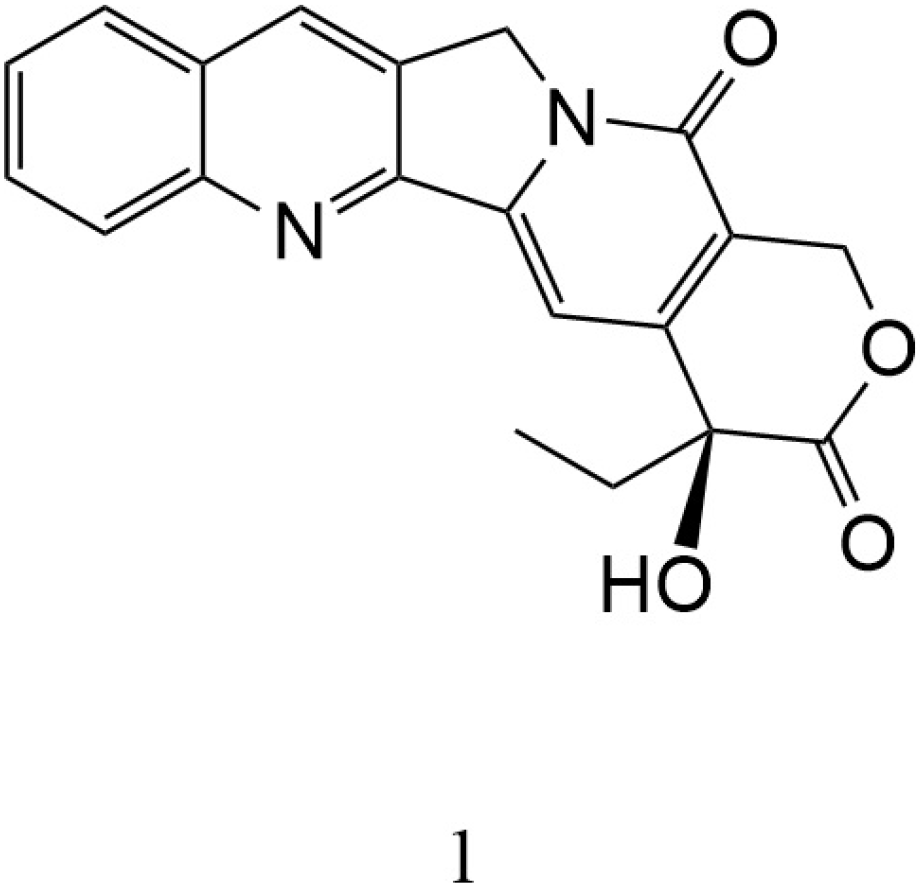

## RESULTS AND DISCUSSION

### Isolation of camptothecin yielding endophytes from *Nothapodytes nimmoniana*

Endophytes residing in the plants may vary with the type of plant tissue.^12,13^ Therefore, leaves, petioles, stem and bark regions from *Nothapodytes nimmoniana* (J.Graham) Mabb. were used as explants for the comprehensive isolation of endophytes. Endophytes emerged from the cut ends of the explants after 5-7 days of incubation in potato dextrose agar medium (HiMedia, Mumbai). A total of 132 visually distinct endophytic strains were isolated from the explants of *N. nimmoniana*.

These isolated endophytes were then individually screened for their ability to produce camptothecin in suspension. Suspension cultures were initiated in duplicate and the biomass was harvested and dried for extraction of camptothecin for quantitative analysis using high performance liquid chromatography (HPLC). Among the 132 different endophytic isolates, 94 produced camptothecin in culture, with yields ranging from 0.1 – 400 μg/g DW (dry weight of biomass).

Four of the highest yielding endophytes were selected for further analysis and their camptothecin yield was confirmed again in freshly initiated cultures (in triplicate). Camptothecin yields obtained from the suspension cultures of these strains are shown in Table 2. Strain P4-6-PE2, isolated from the petiole region of a young plant gave the highest yield of camptothecin in suspension (426.7±33.6 μg/g), followed by strain P4-4-LE2 isolated from the leaf region of the plant, with a yield of 403.3±41.6 μg/g. It is to be noted that the yield of camptothecin obtained from the natural plant of *C. acuminata* varies from 0.5 – 2.5 mg/g ^14^ and the yield from *N. nimmoniana* ranges from 0.8 – 3 mg/g.^15^ However, wide variations in the plant’s natural yield due to climatic and geographical factors, its slow growth rate and unavailability due to overharvesting have been the key limitations to meet the market demand of camptothecin. On the contrary, a microbial fermentation process, amenable to optimization and scale-up for enhanced yield & productivities, can be an excellent alternative to the natural plant source for large scale sustainable production of camptothecin.

**Table 2.** Highest yielding endophytes with their camptothecin yield, accession and deposition details

| S.No | Explant | Endophytes | Dried biomass<br>obtained from<br>suspension<br>cultures<br>(g/l) | Camptothecin<br>yield<br>(µg/g) | Species full<br>name | Genbank<br>accession<br>number | NCIM<br>catalogue<br>number |
| --- | --- | --- | --- | --- | --- | --- | --- |
| 1 | Petiole | P4-6-PE2 | 6.24 ± 0.09 | 426.7 ± 33.6 | *Alternaria*<br>*alstroemeriae* | MN795638 | NCIM1408 |
| 2 | Leaf | P4-4-LE2 | 8.94 ± 0.13 | 403.3 ± 41.6 | *Alternaria*<br>*burnsii* | MN795639 | NCIM1409 |
| 3 | Leaf | P4-1-LE1 | 8.52 ± 0.61 | 269.4 ± 53.9 | *Alternaria*<br>*alstroemeriae* | MN795640 | NCIM1441 |
| 4 | Leaf | P5-4-LE1 | 9.78 ± 0.23 | 62.5 ± 0.6 | *Alternaria*<br>*angustiovoidea* | MN795641 | NCIM1442 |

Camptothecin production was confirmed in the suspension cultures of these camptothecin yielding endophytic strains using tandem mass spectrometry (LC-MS/MS). As expected, a precursor mass to charge ratio (m/z) of 349.12 corresponding to that of camptothecin was observed in the authentic standard and in the extracts from all the shortlisted (top 4) endophytic strains (Figure 1). Further, for structural confirmation, the precursor m/z (349.12) was fragmented by applying collision energy of 35 electron Volts (eV) and an m/z of 305.13 (fragment) after the loss of CO_2_ was witnessed (Figure S1, Supporting information), confirming the presence of camptothecin in the extract sample.

**Figure 1.**
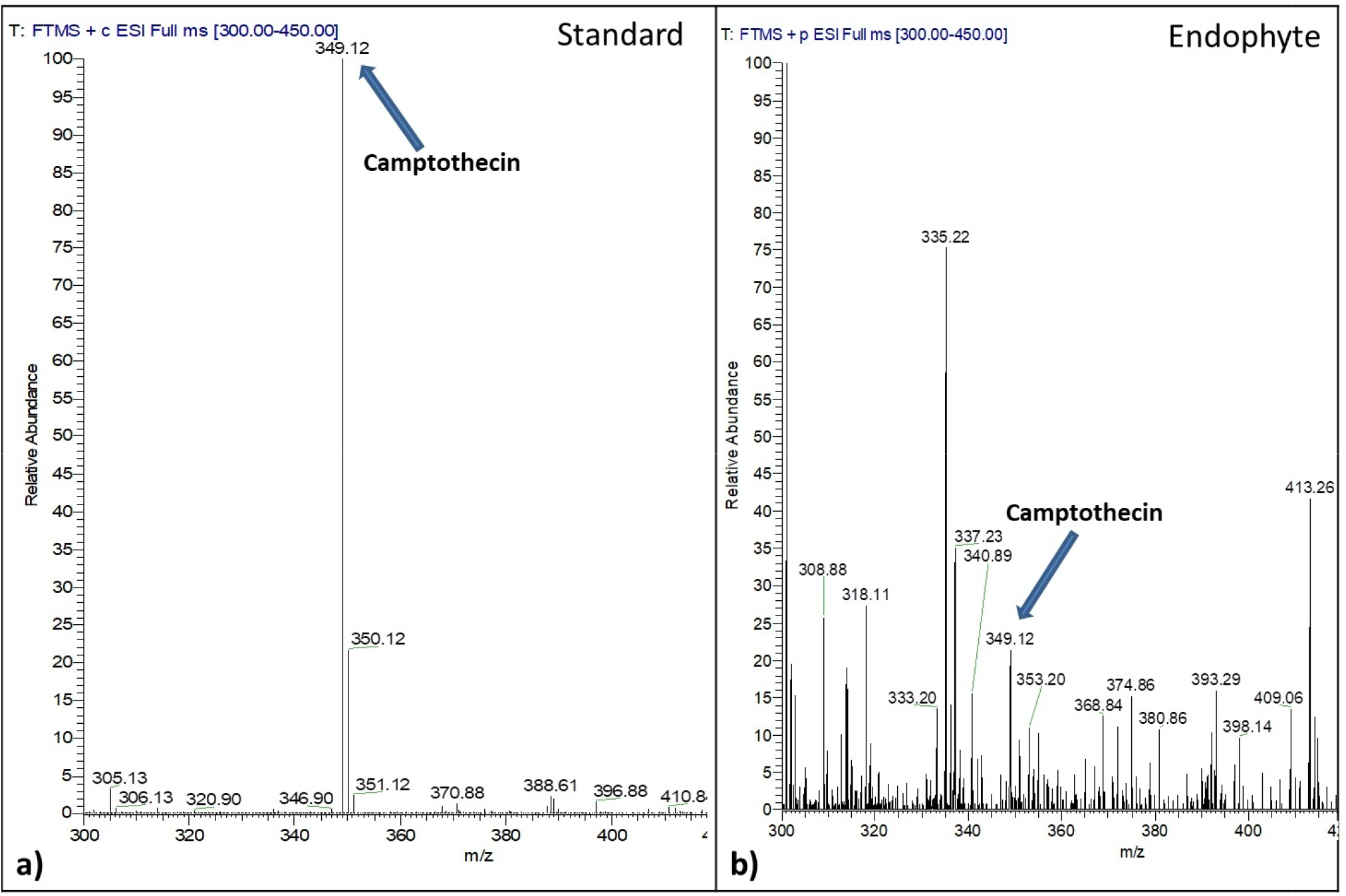
Identification of camptothecin from the endophytic extract using mass spectrometry. An m/z of 349.12 corresponding to camptothecin visualized in both the standard (a) and the isolated endophyte (b)

The presence of camptothecin in the endophytic extracts was confirmed using thin layer chromatography (TLC). Along with the camptothecin producing culture extract, a non-camptothecin producing culture extract was also spotted on the TLC plate for analysis. As shown in Figure S2, supporting information, all four camptothecin yielding endophytic extracts gave spots corresponding to that of authentic camptothecin. In contrast, the spot was absent from a non-camptothecin producing strain.

### Molecular characterization of the selected high camptothecin yielding endophytes

Endophytic strains grown on plates made with PDA medium showed difference in their morphological appearance and all of them formed spores when viewed under a microscope (Figure S3, Supporting information). All four isolates were found to be different species of *Alternaria* genus. Among the four, the highest yielding strain was found to be *A. alstroemeriae* with a yield of 426.7±33.6 μg/g and the second highest was found to be *A. burnsii* with a yield of 403.3±41.6 μg/g. Accession numbers from NCBI Genbank and the respective catalogue numbers of each strain deposited in NCIM have been listed in Table 2.

#### Sustainable production of camptothecin in the highest yielding endophyte

Sustainable production of camptothecin in the two highest camptothecin yielding endophytes was assessed in suspension culture generated from the 1^st^ through the 12^th^ generation slants. The highest yielding strain (*A. alstroemeriae* NCIM1408) demonstrated a sharp decrease in the camptothecin yield from 426.7±33.6 μg/g DW in culture developed from its 1^st^ generation slant to 17.9±0.7 μg/g DW in culture from its 12^th^ generation slant. This result was in accordance with the well-known product yield attenuation observed in axenic cultures of endophytes (Table 1). Interestingly, unlike the highest yielding strain, the second-highest yielding strain (*A. burnsii* NCIM1409) could demonstrate sustainable production of camptothecin (up to ~200 μg/g) in culture even from its 12^th^ generation slant used as inoculum. As shown in Figure 2, after an initial decrease in the yield to 254.1±17.9 μg/g DW in cultures developed from the second generation slant, this endophyte demonstrated a steady yield of ~200 μg/g DW in cultures from 3^rd^ till 12^th^ generation slant.

**Figure 2.**
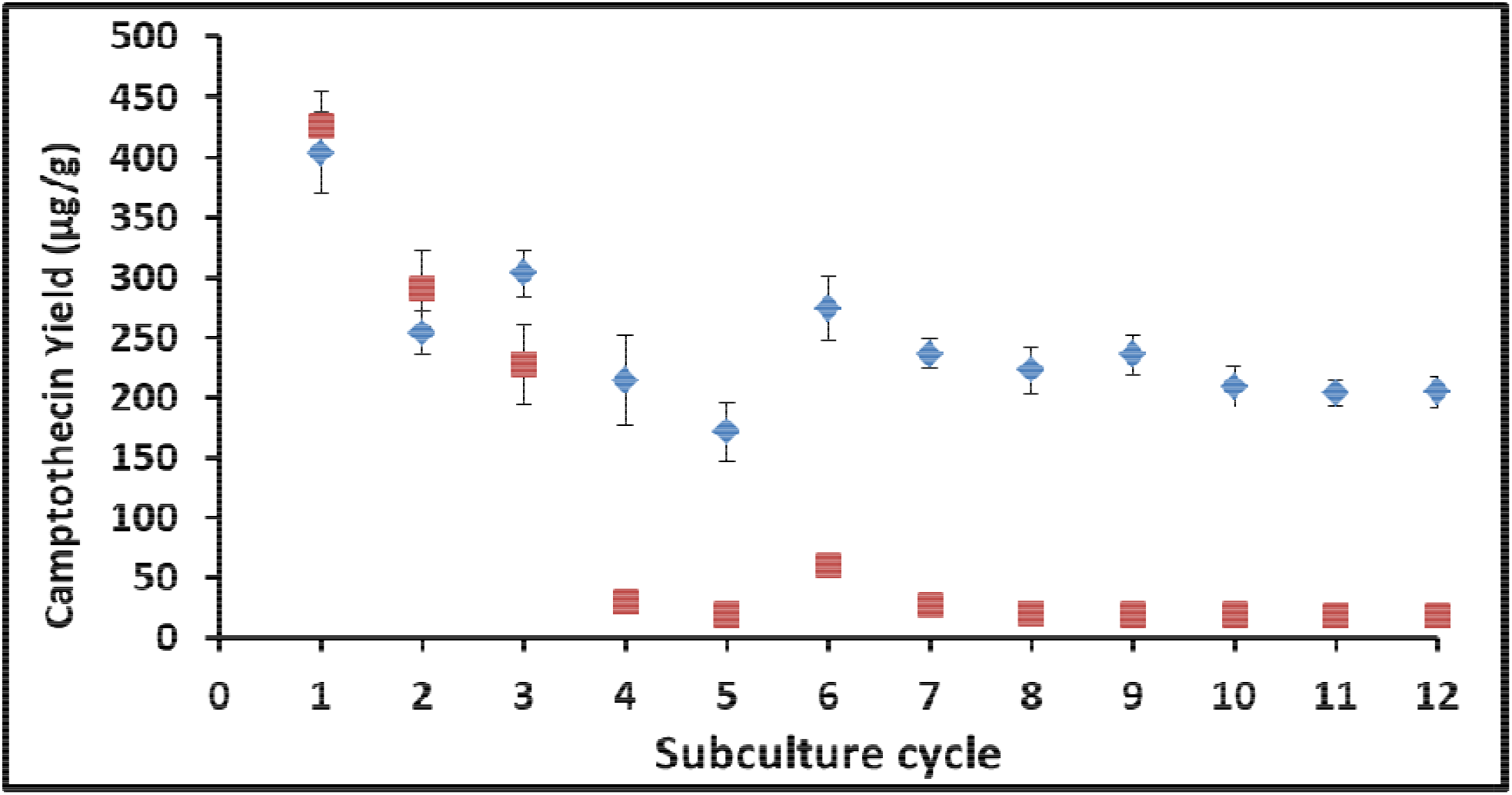
Stability analysis on camptothecin yield (Average yield ± SEM) from the two high yielding endophytic strains (♦ *A. burnsii* NCIM1409 & ■ *A. alstroemeriae* NCIM1408)

These results demonstrate that the highest yielding strain may not always be a sustainable producer, and it is equally important to consider sustainable production along with product yield of the metabolite during the screening and selection of a potential endophyte for successful bioprocess development.

Though numerous endophytes are present in the plant, only a few of the isolated strains have the potential to produce the host-specific metabolite and many have shown attenuation upon subculturing.^9,16^ Various factors were stated to play a role in the endophyte’s non-sustainability, including their evolutionary relationship with the host plant,^17^ and lack of host stimuli ^18^ or gene silencing under axenic conditions.^19^ Thus, comprehensive bioprospecting and screening are required to identify a sustainable and high product yielding strain of endophyte.

### Confirmation of camptothecin biosynthesis by *A. burnsii* via carbon metabolism

Existence of host independent biosynthetic machinery in endophytes has been doubted due to product yield attenuation under an axenic state.^20^ To confirm the endophytic strain has the endogenous ability to synthesize camptothecin, the cultures were fed with D-[U-^13^C]-glucose and ^13^C label incorporation into camptothecin was monitored. The fungal cultures fed with D-glucose and D-[U-^13^C]-glucose showed similar growth, evident from the biomass (~8 g/l DW) achieved after 8 days of cultivation carried out under similar conditions. An m/z of 349.12 corresponds to that of camptothecin with all unlabelled carbon atoms and the presence of one ^13^C labeled carbon would increase the molecular weight of camptothecin by 1 dalton. Since camptothecin has 20 carbons, there is a possibility that 1-20 carbon atoms in the camptothecin molecule produced can get labelled in the culture extract when grown on D-[U-^13^C]-glucose. This was verified by tandem mass spectrometry confirming the increase in the molecular mass of camptothecin corresponding to the number of carbon atoms labelled in it. The mass spectrum revealed majority of carbon atoms (2-20) getting labelled in camptothecin. This was further supported by the MS-MS fragmentation pattern of the most abundantly labelled camptothecin molecules in the sample after the expected removal of labelled and unlabelled CO_2_ (Table S1 and Figure S4, Supporting information). Moreover, the absence of unlabelled camptothecin mass peak in D-[U-^13^C]-glucose fed sample (Figure 3b) and absence of labelled camptothecin mass peak in D-glucose fed sample (Figure 3a & 3c) could also substantiate labelled glucose metabolism and its incorporation in the synthesis of camptothecin by the organism.

**Figure 3.**
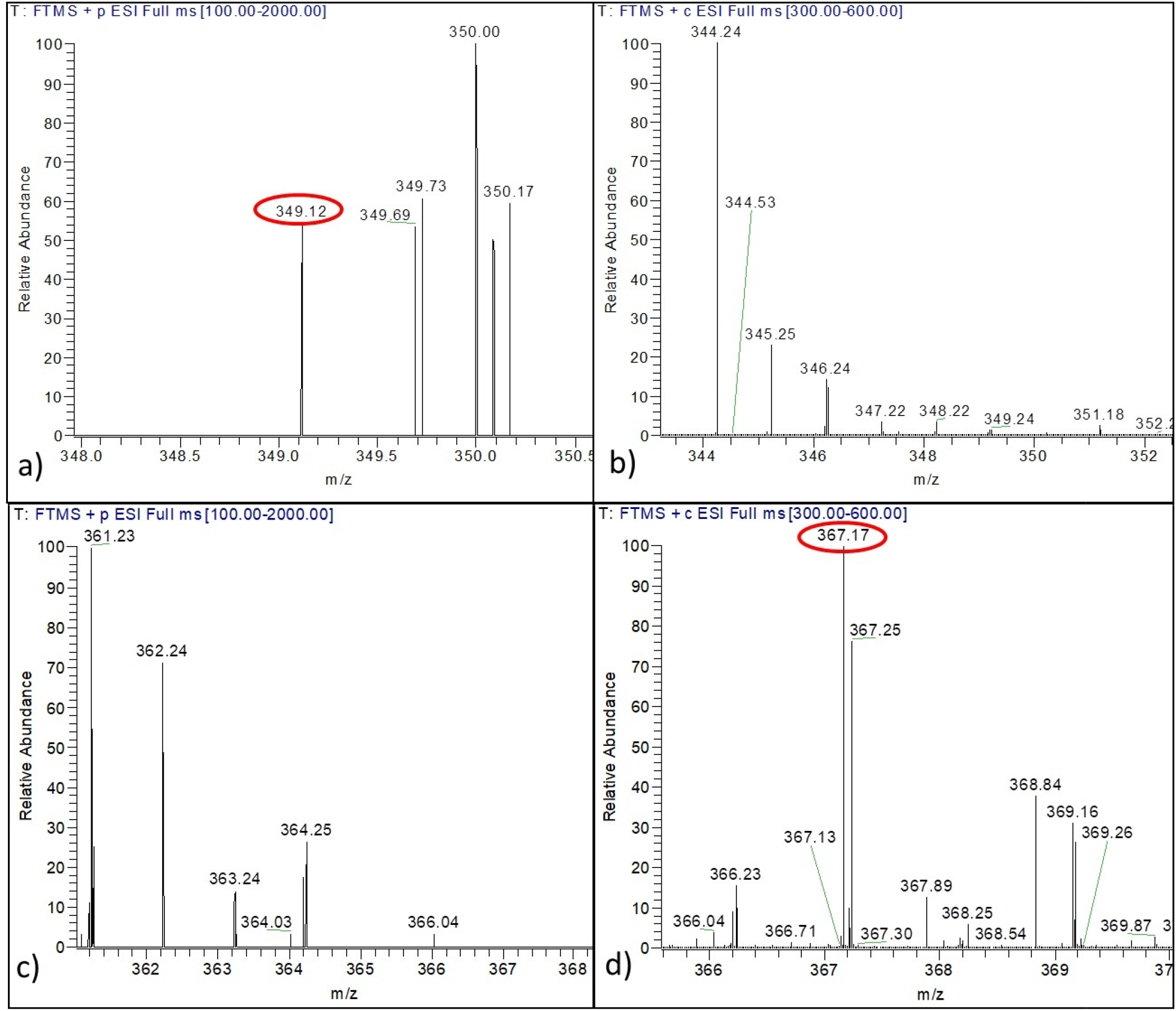
Confirmation of camptothecin biosynthesis in *A. burnsii* using D-[U-^13^C]-glucose. Mass spectrum of fungal extract shows that non labelled camptothecin m/z of 349.12 was visualized only in D-glucose fed samples (a) and m/z corresponding to labelled camptothecin was not present in them (c). In case of D-[U-^13^C]-glucose fed cultures non labelled camptothecin m/z of 349.12 was not visualized (b) whereas an m/z of 367.17 corresponding to 18 carbons in camptothecin being labelled was seen (d).

Interestingly, the mass peak corresponding to camptothecin with 18 labelled carbons (^13^C_18_C_2_H_16_N_2_O_4_) showed highest relative abundance (Figure 3 and S5, Supporting information) while m/z corresponding to 20 labelled carbons (^13^C_20_H_16_N_2_O_4_) in camptothecin was relatively less abundant leading to only its precursor mass (369.18) getting detected and not its MS-MS fragment. This was presumably due to the fact that potato infusion present in the medium is a source of unlabelled organic carbon which could have diluted the ^13^C incorporation from D-[U-^13^C]-glucose. Based on the MS data obtained, this accounts to ~10% dilution with unlabelled carbons, leading to maximum relative abundance of camptothecin with18 labelled carbons (^13^C_18_C_2_H_16_N_2_O_4_). The supportive calculation can be seen in the supplementary section (Table S2, Supporting information).

#### Cytotoxicity analysis of the camptothecin extract from *A. burnsii* on cancer cell lines

As the source of camptothecin was novel, the cytotoxicity analysis was also carried out to further substantiate the potential commercial utilization of the microbial strain for various anti-cancer applications. The cytotoxicity of the solvent extract of camptothecin (~25% pure) from the biomass of *A. burnsii* was tested on various cancer cell lines which includes human breast cancer (MCF7), human ovarian adenocarcinoma (SKOV3), human non-small cell lung carcinoma (H1299), colon adenocarcinoma (Caco-2, HT29), and non-cancerous embryonic kidney (HEK293T) cell lines. The IC_50_ values determined for the camptothecin extract and standard camptothecin against all the cancer cell lines studied have been shown in Table 3 as calculated from the plot of % cell viability vs. log concentration of the extract (Figure S6, Supporting information). Apart from MCF7, all other cancer cell lines showed a lower IC_50_ value for camptothecin extract from *A. burnsii* than the standard camptothecin (≥90 % pure), which could be due to the effect of other metabolites present in the extract along with camptothecin (Table 3). Among all the cancer cell lines used, camptothecin extract from *A. burnsii* was found more toxic toward lung and colon cancer cells which is in accordance with the reports on derivatives of camptothecin.^4^

**Table 3:** Cytotoxic activity of camptothecin extract from *A. burnsii* and camptothecin standard on the various cancerous and non-cancerous cell line

| Tissue | Cell Line | IC<sub>50</sub> value of camptothecin in the extract of *A. burnsii* (nM) | IC<sub>50</sub> value of camptothecin standard (nM) |
| --- | --- | --- | --- |
| Breast | MCF7 | 44.6 ± 12.6** | 14.7 ± 0.7 |
| Lung | H1299 | 13.6 ± 1.7** | 93.3 ± 22.6 |
| Ovary | SKOV3 | 117.4 ± 10.4* | 165.9 ± 31.2 |
| Colon | HT29 | 50.1 ± 11.2*** | 467.4 ± 90 |
| Colon | Caco-2 | 26.9 ± 3.5*** | 70.7 ± 7.7 |
| Non-cancerous embryonic kidney | HEK293T | 281.5 ± 33* | 158.4 ± 55.6 |

Confidence interval for statistically different IC<sub>50</sub> of the camptothecin extract, in comparison to that of the standard, for each cancerous cell line: \* $P \leq 0.05$ ; \*\* $P \leq 0.01$ ; \*\*\* $P \leq 0.001$

The cytotoxicity of the camptothecin extract from *A. burnsii* was further supported by clonogenic (Figure 4) and wound healing assay (Figure 5), where colony formation and healing were inhibited on the selected SKOV3 cell line.

**Figure 4.**
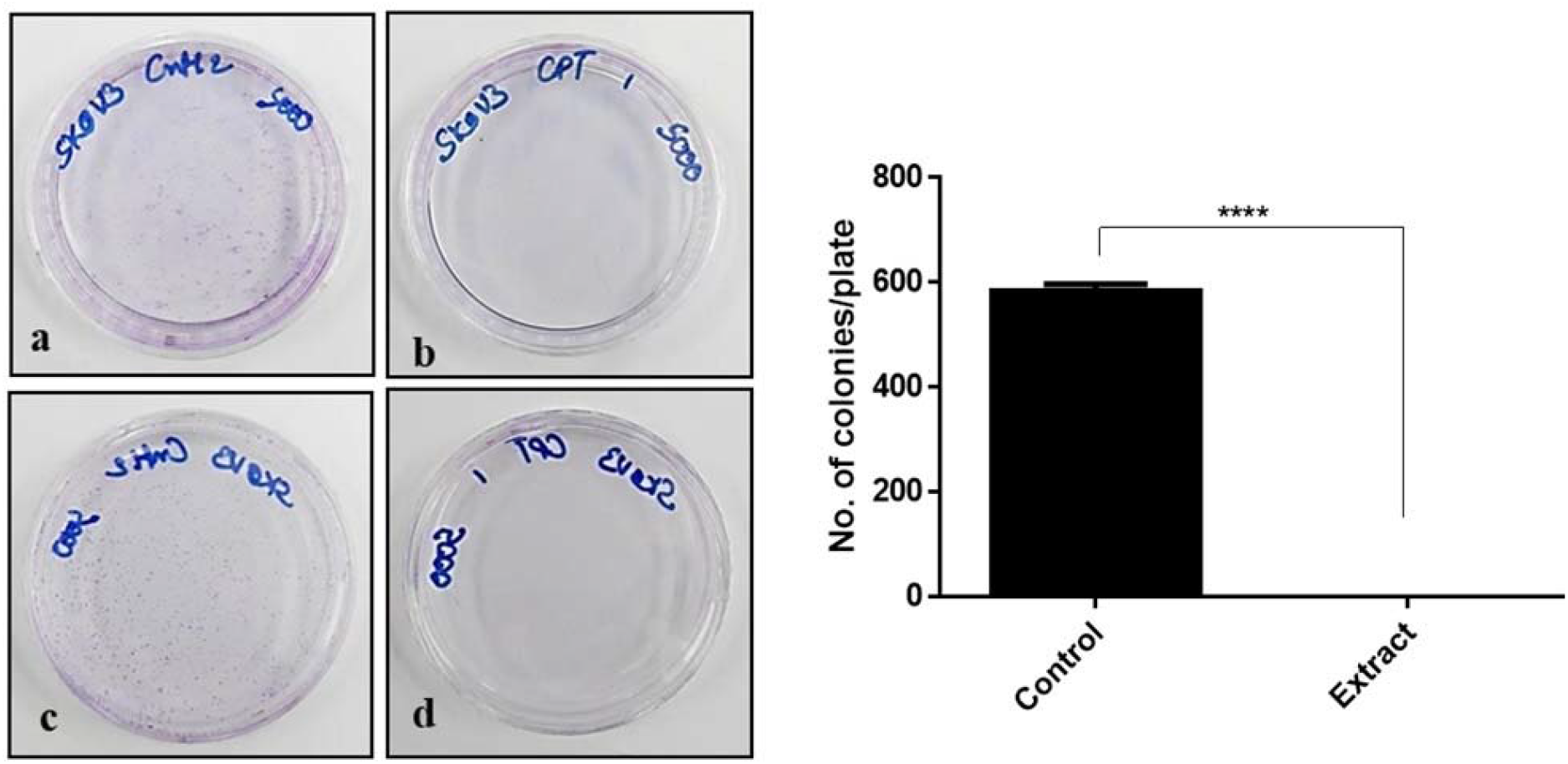
Complete inhibition of colony formation in the SKOV3 cell line by *A. burnsii* extract at IC_25_. SKOV3 control plates showed colony formation (a, c), whereas the treated plates showed no colony formation (b, d).

**Figure 5.**
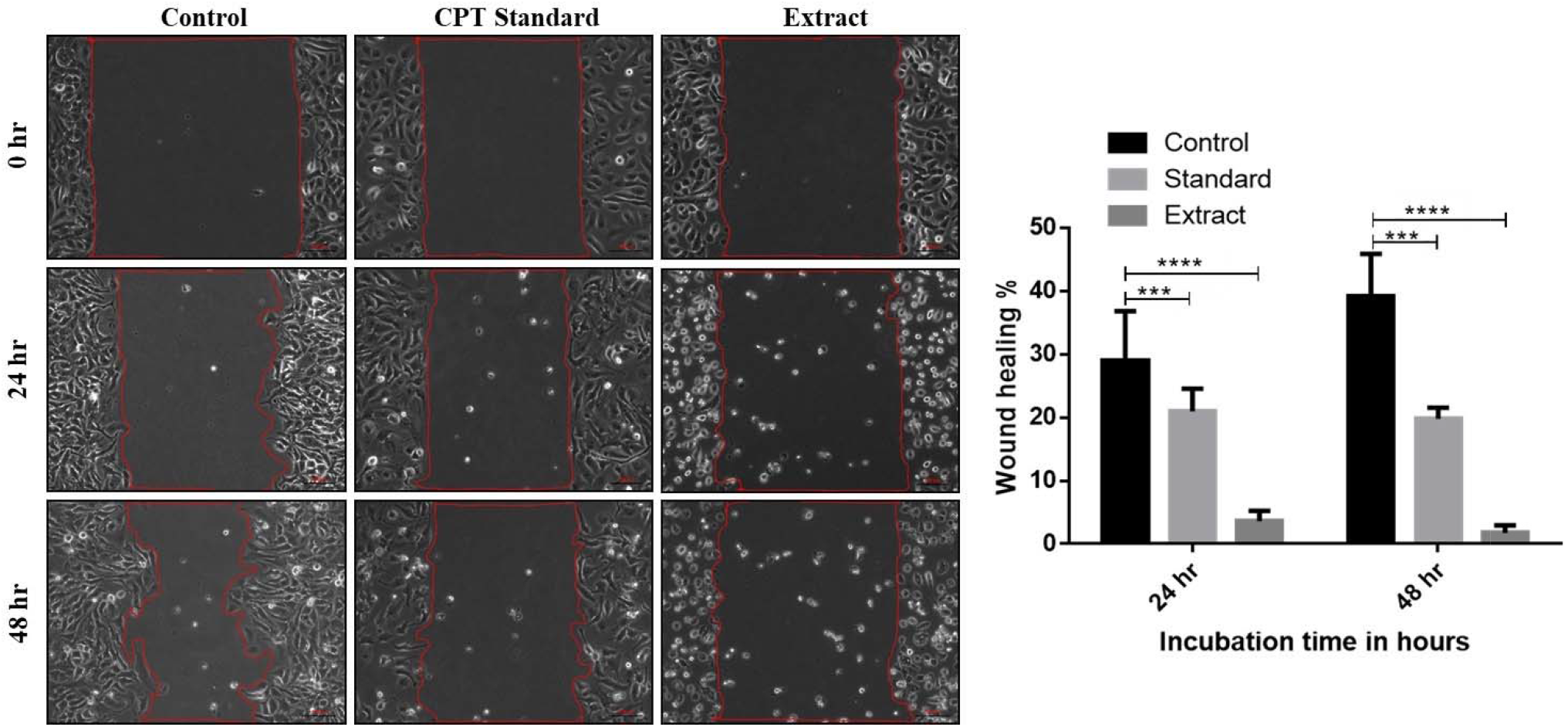
Minimal wound healing observed with *A. burnsii* extract-treated SKOV3 cell line. Control cells showed 29% and 39% healing after 24 and 48 h respectively, while the standard camptothecin treated cells showed 20% and 19% healing after 24 and 48h respectively. The crude extract-treated cells showed only 1% healing after 48 h.

In conclusion, a high camptothecin yielding fungal endophyte *A. burnsii* NCIM1409 was isolated from *N. nimmoniana* which could demonstrate sustainable production of camptothecin under axenic conditions. Production of camptothecin by the microbial culture was verified via ^13^C labeled glucose metabolism leading to labeled camptothecin biosynthesis. Interestingly, the crude camptothecin extract from the fungi demonstrated better cytotoxic activity (lower IC50) than the standard camptothecin (>90 % purity) against the tested lung, colon and ovarian cancer cell lines and relatively less cytotoxicity against the non-cancerous cell line. The isolated strain can be used to develop a microbial-based sustainable bioprocess for large scale in vitro production of camptothecin for anti-cancer applications as an alternative to natural plant extraction.

## EXPERIMENTAL SECTION

### General experimental procedures

Endophytes were isolated from 6 different plants of *N. nimmoniana* cultivated in the University of Agricultural Science, Bengaluru campus (13.0762 °N, 77.5753 °E). The plant specimen was authenticated as *N. nimmoniana* (J.Graham) Mabb. of the family Icacinaceae and deposited in Foundation for Revitalisation of Local Health Traditions (FRLH) herbarium, Bengaluru (FRLH 122013). The collected plant material (leaves, petioles, stem and bark) from young and mature plants was washed thoroughly in running tap water to remove the dust particles and then with sterile water. Surface sterilization was carried out within 10 min of excising the explants from the parent plants since the time between excision of the explant and isolation can reduce the viability of endophytes residing in the plant parts.^21^ For surface sterilization of the explants, leaves and petioles were treated with 1 % (v/v) sodium hypochlorite for 1 min and then washed with sterile distilled water to remove sodium hypochlorite from the plant parts. The washed explants were then treated with 70 % (v/v) ethanol for 1 min and then washed again thoroughly with sterile distilled water to remove residual ethanol, if any, from the plant parts. A similar procedure was carried out for the stem and bark regions, except that the treatment with 1 % (v/v) sodium hypochlorite was increased to 3 min and that with 70 % (v/v) ethanol was up to 5 min. The sterile explants were then cut into small pieces (~ 1 cm^2^) and placed on the Petri dishes having PDA medium (HiMedia, Mumbai) for incubation at 28 °C in a Biological Oxygen Demand (BOD) chamber for 7 days to allow the endophytes to emerge and grow from the cut ends. Morphologically distinct endophytes emerged from the cut ends after 5-7 days of incubation and were subsequently removed and plated separately on a fresh medium to obtain pure colonies of the endophytes.

A loop full of the mycelia was streaked on slants made with 5 ml of potato dextrose agar medium (HiMedia, Mumbai) and incubated at 28 °C for 7 days with an initial pH of 5.6. The slants were washed using 5 ml of saline (0.9 % (w/v) NaCl) and were used as inoculum (2 % v/v) for initiating suspension culture. Suspension cultures (in duplicate) were established with 50 ml of potato dextrose broth (HiMedia, Mumbai) in 250 ml Erlenmeyer flasks at an initial pH of 5.6. The suspension culture was allowed to grow in an incubator shaker at 28 °C and 120 rpm. The shake flasks (in duplicate) were harvested after 8 days of the cultivation period for the estimation of biomass (g/l, DW) and camptothecin yield (μg/g DW) as per the protocol reported elsewhere.^22,23^

### Biomass estimation

The biomass was harvested by centrifuging the cell suspension at 12,857 x g for 15 min and discarding the supernatant. The pelleted cells were then washed with distilled water to remove traces of medium components and again separated by centrifugation at 12,857 x g for 15 min. Washed biomass was then dried in a hot air oven at 60 °C by spreading it on pre-weighed glass Petri plates until constant dry weight was achieved.

### Extraction of camptothecin

Camptothecin extraction from the fungal biomass was done as per the protocol reported earlier.^23^ Briefly, the dried biomass (0.3 g) was dissolved in 20 ml of distilled water and homogenized using a mortar and pestle, followed by liquid – liquid extraction, repeated thrice using 50 ml of chloroform: methanol solvent mixture (4:1). The organic layer with extracted camptothecin was then collected and evaporated using a rotary evaporator. The dried camptothecin extract was re-suspended in 1 ml of DMSO: methanol (1:50) and filtered through a 0.2 μm filter for further analysis.

### Quantification of camptothecin by HPLC

Twenty microliters of the camptothecin extract was injected in a reverse phase HPLC (LC-20AD, Shimadzu, Japan) at a flow rate of 0.8 ml/ min using 25 % acetonitrile as the mobile phase. ODS Hypersil gold column (Thermo Scientific) with a particle size of 5 μm was used as the stationary phase at a column temperature of 30 °C. The absorbance of camptothecin was measured at 254 nm by a photodiode array detector.^24^ Concentration of camptothecin in the sample was estimated using a standard calibration curve of peak area vs. known concentration of camptothecin (Figure S7, Supporting information), generated using authentic samples of camptothecin (>90 % purity, Sigma Aldrich, MO, USA). The retention time of camptothecin from the standard was found to be 19.8 min and the area of the peak obtained from the endophytic extracts at the same retention time (Figure S8, Supporting information) was compared with the standard plot of camptothecin and the corresponding concentration and yield from the different fungal isolates were calculated.

### Identification of camptothecin by LC-MS/MS

Camptothecin production in suspension culture of the high yielding isolates was further confirmed using tandem mass spectroscopy. This was based on qualitative identification of camptothecin, extracted from the harvested biomass, using an orbitrap elite mass spectrometer (ThermoScientific, Breman, Germany). A reverse-phase column (ODS Hypersil C_18_ column, 256×4.5 mm, 5 μ) was connected to a heated electrospray ionization (H-ESI) source with the heater temperature maintained at 325 °C. The sheath gas flow rate was set at 40 arb, aux gas flow rate at 20 arb and sweep gas flow rate at 2 arb. The spray voltage for the H-ESI source was set at 4.5 kV and the capillary temperature was maintained at 300 °C. MS grade water with 0.01 % formic acid (solvent A) and acetonitrile (solvent B) in the ratio 3:1 was used as mobile phase at a flow rate of 500 μl/min. Samples were injected into a 20 μl sample loop and analyzed for 60 mins in the LC coupled with mass spectrometry set up in a mass scanning range of 300-450 m/z. The precursor ions were subjected to collision induced dissociation (CID) of 35 eV to obtain fragment ions. Camptothecin, with a molecular formula of C_20_H_16_N_2_O_4_, when analyzed in LC-MS under positive mode, yielded an intact mass [M+H] of 349.12. Further, for structural confirmation, by applying CID of 35 eV, the precursor m/z (349.12) could be fragmented to an m/z of 305.13 with a loss of CO_2_.^3^

### Qualitative analysis of camptothecin by TLC

Presence of camptothecin in the endophytic extracts can also be identified using TLC. TLC was performed using silica gel 60 F_254_ aluminium plates (Merck, Germany). Extracts of camptothecin (10 μl) from selected endophytes were spotted on the TLC plate with the standard camptothecin sample used as positive control. Chloroform-methanol (20:1) solvent system was used as the mobile phase. The plates were placed in the mobile phase in a pre-saturated chamber and the samples were allowed to be drawn upward on the TLC plates via capillary action for the separation of the sample components. As camptothecin is known to exhibit fluorescence under UV light, the TLC plates were dried and visualized at 254 nm in a UV chamber as per the protocol reported elsewhere. ^25,26^

### Molecular characterization of the selected endophytes

Freshly grown mycelia (200 mg) from the suspension cultures of the selected endophytes were taken and washed with ethanol before grinding to a fine powder with liquid nitrogen and sterile sea-sand in a mortar and pestle. The homogenized sample was then transferred into a sterile 1.5 ml microcentrifuge tube and 200 μl of cell-lysis buffer (Nucleo-pore DNA Sure Plant Mini Kit, Genetix Biotech, India) was added. This mixture was incubated at 70 °C for 10 min and then chloroform was added (100 μl) and mixed thoroughly. Phase separation was then carried out in the microfuge tube after centrifugation at 20000 g for 15 min. The top aqueous layer was separated into a sterile 1.5 ml microcentrifuge tube for the isolation of DNA using nucleopore DNA sure plant mini-kit as per manufacturer’s protocol (Genetix Biotech, India). Genomic DNA was isolated from the four highest yielding endophytic strains.

Isolated genomic DNA was amplified using internal transcribed spacer 1 (ITS1) (5′ TCCGTAGGTGAACCTTGCGG 3′) and internal transcribed spacer 4 (ITS4) (5′ TCCTCCGCTTATTGATATGC 3′) primers,^27^ along with the Phusion High Fidelity DNA polymerase (New England Biolabs, Ipswich). Touch-down method of PCR was used for increased specificity of primer amplification in a Surecycler 8800 (Agilent Technologies, Santa Clara, CA). The upper limit of annealing temperature was set at 65 °C and the lower limit at 53 °C. The PCR conditions used were as follows: initial denaturation for 5 min at 95 °C, 35 cycles of amplification including (i) denaturation at 95 °C for 1 min, (ii) primer annealing to DNA at 65 – 53 °C for 45 s, (iii) primer extension at 72 °C for 2 min along with a final extension for 10 min at 72 °C. The purified polymerase chain reaction (PCR) products showed bands between 500-700 bp corresponding to that of the conserved ITS regions in fungi. The amplified products were purified using a Qiaquick PCR purification kit as per manufacture’s protocol (Qiagen, California, USA) and the size was confirmed using agarose gel electrophoresis. In the gel made with 0.8 % (w/v) agarose, 10 μl of the amplified product and 2 μl of 6X gel loading dye were added in the wells and the system was run at 50 V for 30 mins before visualization of the amplicons on the gel under a UV transilluminator (GelDoc, BioRad, Italy). The purified PCR products were then sequenced by Sanger dideoxy method using AB 3130 Genetic Analyzer (Applied Biosystems, Foster City, CA). Forward and reverse sequences were aligned to identify the consensus sequence consisting of ITS1, 5.8S and ITS4. These sequences were compared against the database of sequences using the nucleotide BLAST algorithm provided by (National Center for Biotechnology Information) NCBI. Also a phylogenetic tree (Figure S9, Supporting information) was constructed using maximum likelihood method based on Jukes-Cantor model ^28^ and the analysis was conducted in MEGA7.^29^ To increase the level of confidence in the taxonomic identifications, the comparisons were restricted to the sequences from the type material by checking the “sequences from type material” box in the general BLAST page.^30^ The search results that displayed the highest query coverage and maximum score were used to determine the identity of the isolated high yielding endophytes. The sequences were deposited in NCBI and the strains were deposited in the National Collection of Industrial Microorganisms (NCIM) Pune, India

### Investigation of sustainable production of camptothecin in the endophytes

Sustainable production of camptothecin was investigated in the 2 highest yielding endophytes isolated in the study. Freshly isolated strains (stored as 25 % glycerol stocks) were streaked separately as slants made of potato dextrose agar medium (5 ml) (HiMedia, Mumbai) and incubated at 28 °C for 7 days with an initial pH of 5.6. These culture slants were used as inoculum for generating suspension and were considered as the first generation (subculture cycle) slants in the study. After 7 days, single mycelia from the first generation slant was picked and streaked on to the new slants with fresh medium (5 ml of potato dextrose agar) under similar conditions. These were considered as the second generation slants. Similarly, subsequent generation slants (up to 12^th^) were created after every 7 days of subculture on to fresh medium.

Suspension cultures from each generation slants were initiated as described earlier. After 7 days of growth, the slant-wash with 5 ml of saline (0.9 % NaCl) was used as inoculum (2 % v/v) for suspension culture. After 8 days of the cultivation period, the shake-flask cultures were harvested (in triplicate) for the estimation of biomass and camptothecin production.

### Investigation of camptothecin biosynthesis using ^13^C labelled glucose

In order to investigate and confirm camptothecin biosynthesis by the endophyte, suspension cultures were initiated using 20 g/l D-[U-^13^C]-glucose (389374, Sigma Aldrich, MO, USA) as the carbon source in 50 ml of sterilized potato infusion as growth medium in 250 ml Erlenmeyer flasks with 2 % (v/v) of inoculum. The suspension cultures were allowed to grow in an incubator shaker at 28 °C and 120 rpm for 8 days of cultivation period before harvest (in duplicate) for the analysis of ^13^C-labelled camptothecin using LC-MS/MS.

For the preparation of potato infusion, 200 g of fresh potatoes were cleaned, peeled and chopped into smaller pieces and boiled for 15 min in 1000 ml of distilled water at 90 °C. The liquid extract was then filtered through cheesecloth to be used as potato infusion. Extract from the culture grown only in potato infusion was used as a negative control to confirm that no camptothecin production was possible without added glucose in the medium.

### Cytotoxicity analysis of extract on cancer cell lines by MTT assay

The extract of camptothecin from the selected endophyte *A. burnsii* was tested for its cytotoxic effect by performing 3-(4, 5-dimethylthiazol-2-yl)-2, 5-diphenyl tetrazolium bromide (MTT) (Sigma) assay on various cancer cell lines including MCF7, SKOV3, H1299, Caco-2, HT29, and HEK293T. All the cell lines were obtained from American Type Culture Collection (ATCC) and were maintained on Dulbecco`s Modified Eagle Media (DMEM) supplemented with 10 % fetal bovine serum (FBS) except SKOV3, which was maintained on McCoy’s 5A (modified) medium supplemented with 10 % FBS at 37 °C in an atmosphere of 5 % CO_2_.

In a 96 well plate, each well was seeded with 2500 cells in a volume of 100 μl of their respective growth medium and incubated for 24 h after which varying concentration (0.01 μM to 51.2 μM) of the extract of camptothecin and standard camptothecin were added, respectively to the medium. The cells were incubated for 72 h with the drug and then 0.5 mg/ml of MTT reagent in the respective medium was added and incubated for 5 h. Then MTT reagent containing medium was removed and DMSO was added to each well after which the optical density was measured using a 96 well plate reader at a dual-wavelength of 570 and 650 nm. MTT reagent forms formazan crystal within live cells, which then produces purple colour when dissolved in DMSO. The cell viability was calculated by comparing the optical density of the treated wells against the non-treated control wells.

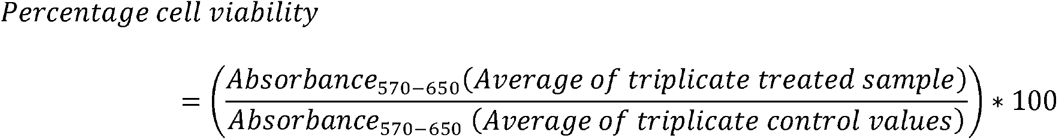

IC_50_ values were calculated using Graphpad Prism 6 (GraphPad Software, San Diego, CA) from the observed cell viability values and SEM.^31^

### Clonogenic and wound healing assay

To investigate the inhibition of the colony formation of these cells in the presence of extract of camptothecin, clonogenic assay was performed. Viable cells (5000 per plate) were seeded and allowed to attach overnight in 60 mm plates. SKOV3 cells were incubated with the extract, at a concentration corresponding to their respective IC_25_ (58.7 nM), for 14 days in medium to facilitate colony formation. The obtained colonies were washed with PBS and fixed using acetic acid: methanol (3:1). The fixed cells were then stained with 0.2 % (w/v) crystal violet. The colonies formed were counted and compared with the DMSO controls in triplicate.

Similarly, a wound-healing assay was performed to study the inhibition of these cells’ migration ability in the presence of the extract of camptothecin. SKOV3 cells were made to form a confluent monolayer on 60 mm plates by seeding 5 million cells on each plate. Using a 200 μL micropipette tip, a wound or scratch was created on the confluent monolayer. The wounded cells were then rinsed with PBS and treated with the extract of camptothecin at a concentration corresponding to their IC_25_ (58.7 nM). The control plates were treated with an equal volume of DMSO. Using phase-contrast microscopy, the wound closure by cell migration was monitored and imaged at 0, 24 and 48 h at the same spot. The wound area was calculated and the percentage of wound healing was compared between the drug-treated cells and DMSO treated cells as control.^31^

## ASSOCIATED CONTENT

### Supporting Information

The supporting information is available free of charge on the ACS Publications website.

MS-MS of camptothecin from the extract (Figure S1), TLC (Figure S2), Microscopic view of the endophytes (Figure S3), Mass spectra of various ^13^C labelled camptothecin (Figure S4), MS-MS of ^13^C labelled camptothecin (Figure S5), Cell viability plots for various cancer cell lines (Figure S6), Standard curve of camptothecin (Figure S7), HPLC chromatograms (Figure S8) and Phylogenetic tree (Figure S9); ^13^C labelled camptothecin masses and their fragments observed (Table S1), Probability of carbons getting labelled in camptothecin (Table S2).

## ACKNOWLEDGEMENTS

The authors thank the Department of Biotechnology, Indian Institute of Technology Madras for the LC-MS facility and Department of Science and Technology (DST), Government of India for the financial assistance for research (Project number: EMR/2015/001418). The authors thank National Cancer Tissue Biobank (NCTB) facility at the Department of Biotechnology, Indian Institute of Technology Madras for, helping with the sequencing of the fungal endophytes.

