## Supporting information for "Sustainable Production of Camptothecin from an *Alternaria* sp. isolated from *Nothapodytes nimmoniana*"

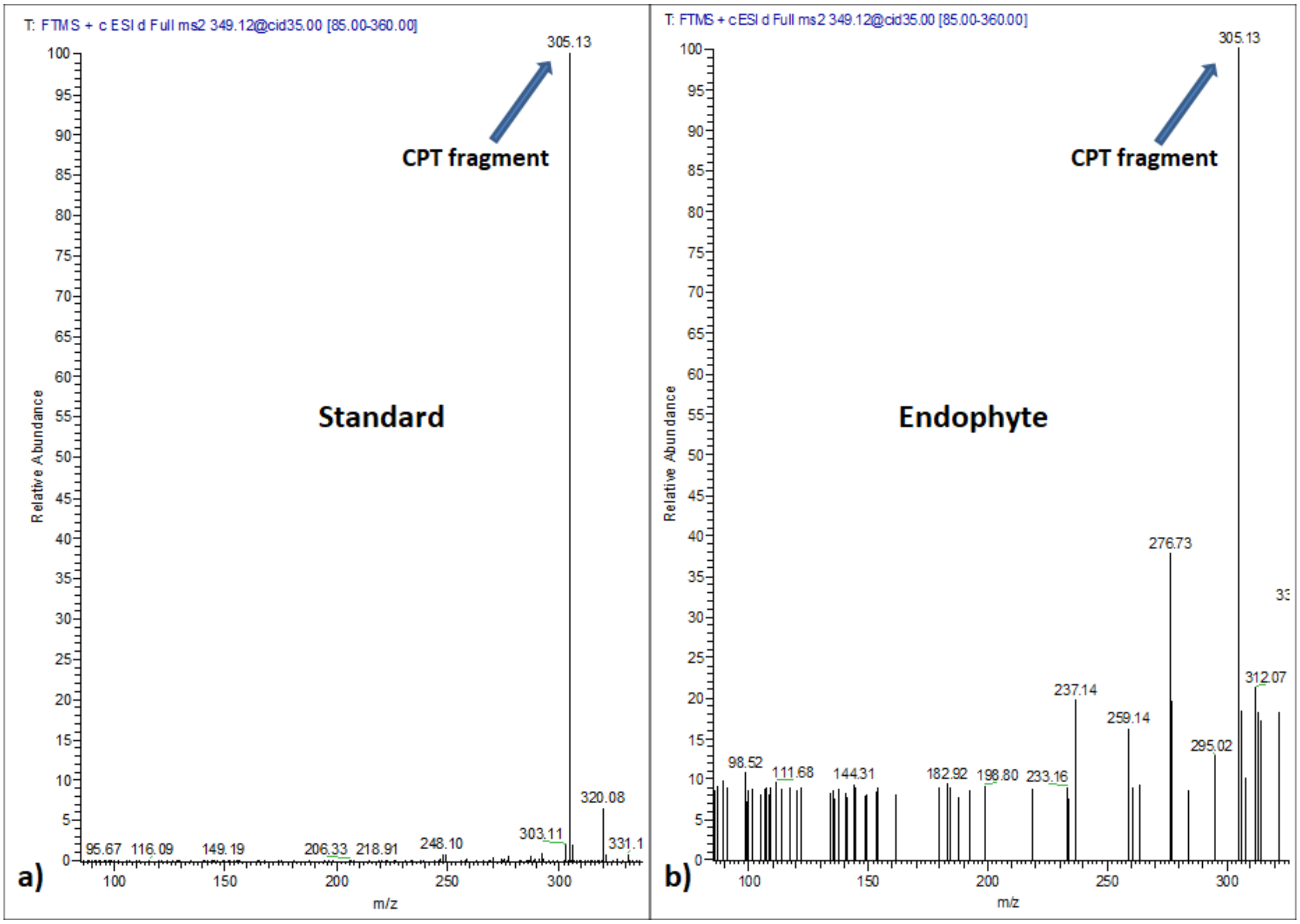


**Figure S1.** Structural confirmation of camptothecin by collision induced dissociation in mass spectrometry. Fragment peak of m/z 305.13 witnessed in both the standard (a) and the isolated endophyte (b).

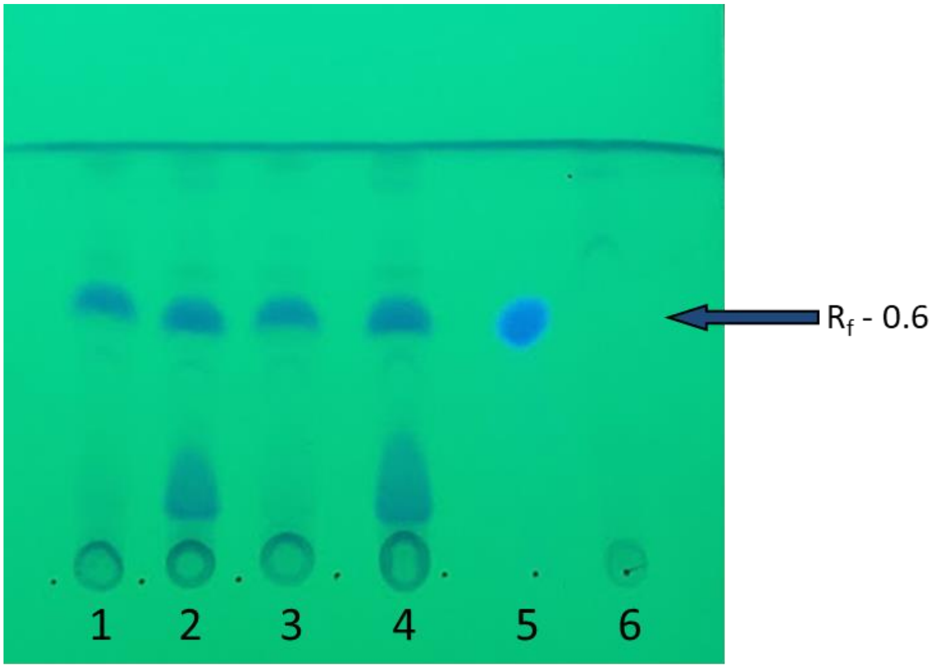


**Figure S2.** Endophytes demonstrating camptothecin presence in TLC plates with an $R_f$ value of 0.6 under 254 nm UV light. Camptothecin producing endophytic extract of P4-6-PE2 (Lane 1), P4-4-LE2 (Lane 2), P4-1-LE1 (Lane 3) and P5-4-LE1 (Lane 4); Standard camptothecin (Lane 5); Non camptothecin producing endophytic extract (Lane 6)

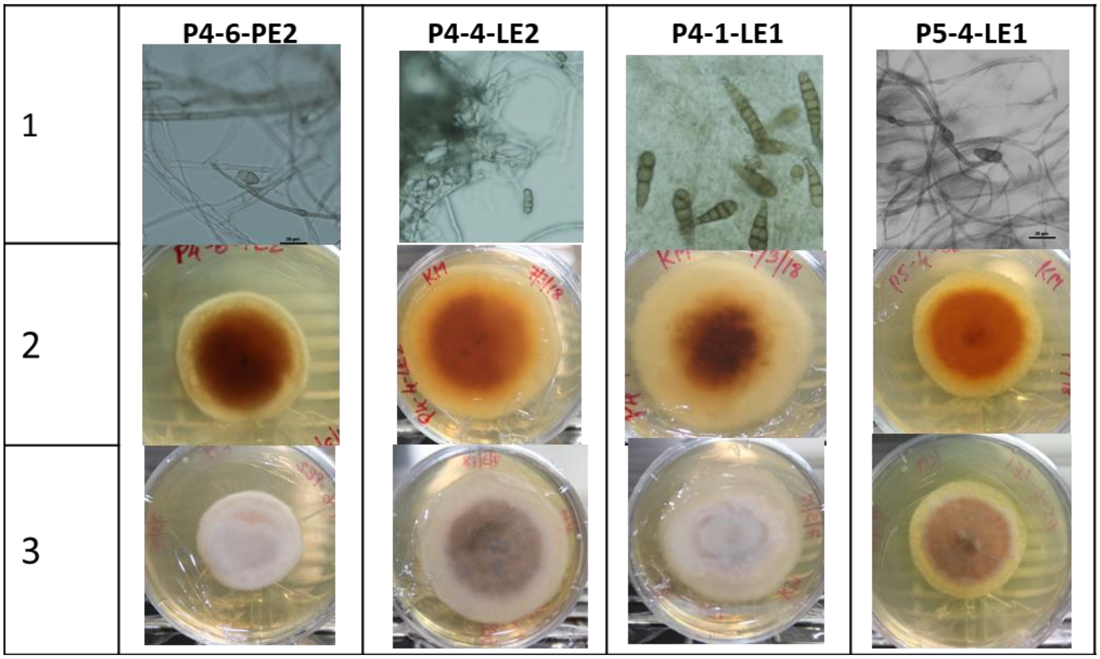


**Figure S3.** Microscopic view of the top 4 camptothecin producing endophytes (Row 1) and morphological appearance of the isolated high camptothecin yielding endophytes from the bottom (Row 2) and top view (Row 3) on plates made with potato dextrose agar (PDA) medium.

**Table S1.** The most abundant masses of $^{13}C$ labelled camptothecin molecules and their corresponding fragment ions after the removal of labelled and unlabeled $CO_2$, that were visualized in the D-[U-$^{13}C$]-glucose fed culture extract.

| Camptothecin with labelled carbons | Corresponding mass peak (m/z) in the spectrum | MS-MS fragment mass peak (m/z) after $CO_2$ removal | MS-MS fragment mass peak (m/z) after $^{13}CO_2$ removal |
|---|---|---|---|
| $^{13}C_9C_{11}H_{16}N_2O_4$ | 358.15 | 314.2 | 313.2 |
| $^{13}C_{10}C_{10}H_{16}N_2O_4$ | 359.15 | 315.2 | 314.2 |
| $^{13}C_{11}C_9H_{16}N_2O_4$ | 360.15 | 316.2 | 315.2 |
| $^{13}C_{13}C_7H_{16}N_2O_4$ | 362.16 | 318.2 | 317.2 |
| $^{13}C_{14}C_6H_{16}N_2O_4$ | 363.16 | 319.2 | 318.2 |
| $^{13}C_{15}C_5H_{16}N_2O_4$ | 364.16 | 320.2 | 319.2 |
| $^{13}C_{16}C_4H_{16}N_2O_4$ | 365.17 | 321.2 | 320.2 |
| $^{13}C_{17}C_3H_{16}N_2O_4$ | 366.17 | 322.2 | 321.2 |
| $^{13}C_{18}C_2H_{16}N_2O_4$ | 367.17 | 323.2 | 322.2 |
| $^{13}C_{19}C_1H_{16}N_2O_4$ | 368.18 | 324.2 | 323.2 |

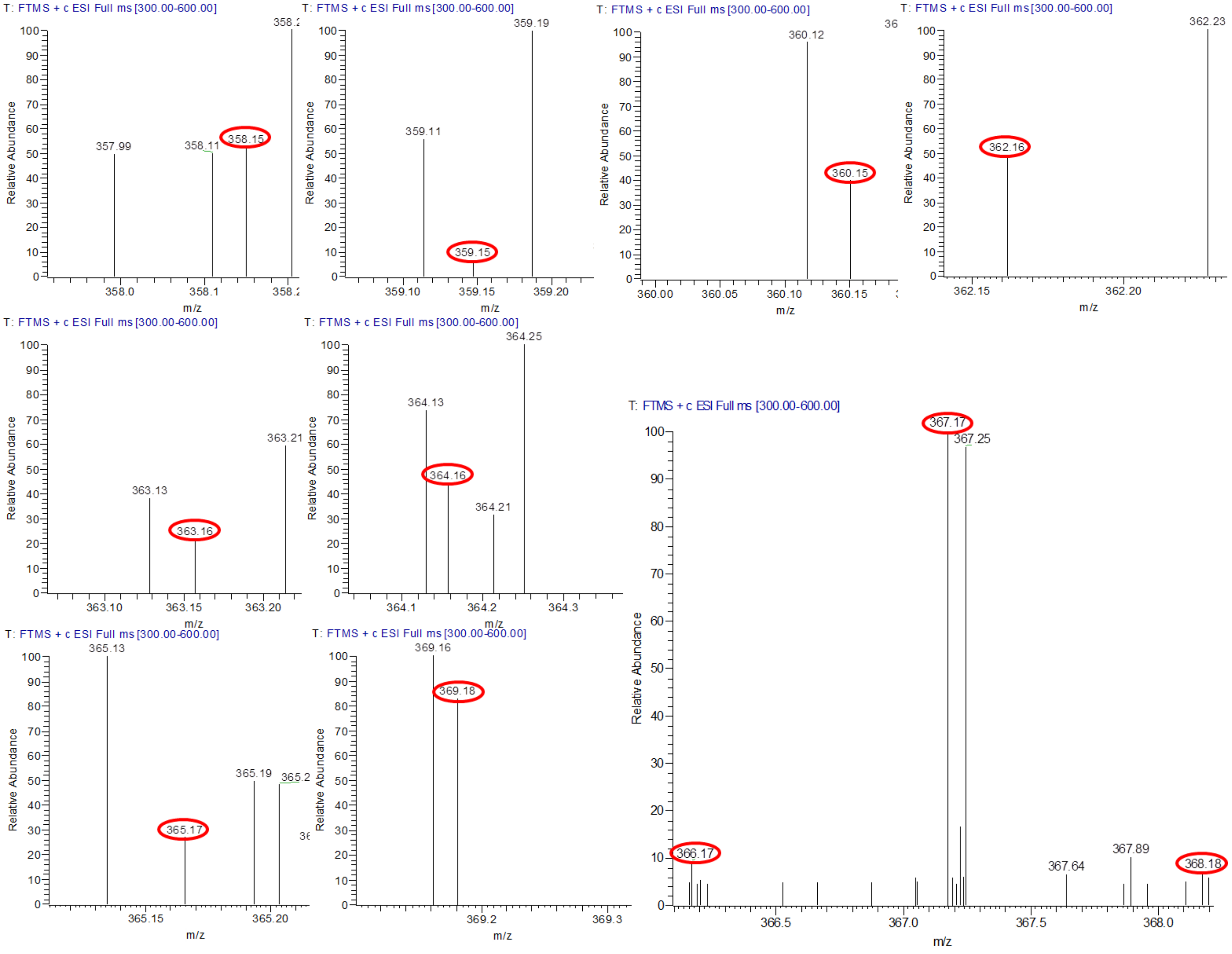


**Figure S4.** Most abundant masses of $^{13}$C labelled camptothecin molecules visualized in the D-[U-$^{13}$C]-glucose fed culture extract

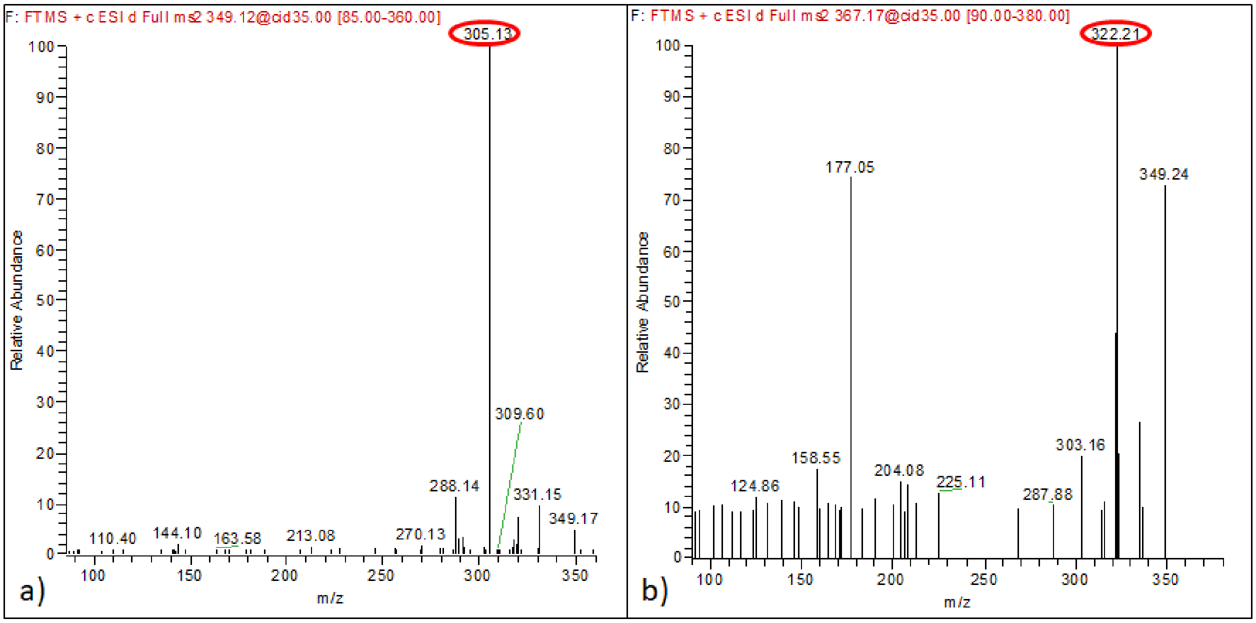


**Figure S5.** Verification of camptothecin biosynthesis in *A. burnsii* at MS2 level in D-[U-$^{13}$C]-glucose study. Fragment peak after removal of $CO_2$ (44 Da) witnessed in the D-glucose fed camptothecin extract (a) and removal of $^{13}CO_2$ (45 Da) was witnessed in D-[U-$^{13}$C]-glucose fed camptothecin extract (b).

**Table S2.** Effect of dilution by the unlabeled carbon on the fraction of labelled camptothecin produced by the culture grown in fully labeled glucose medium supplemented with potato infusion.

| If probability of each carbon getting labelled is X, then probability of camptothecin produced with 'n' number of carbon atoms labelled $(M_n) = X^n (1-X)^{20-n} (^{20}C_n)$ | If the dilution by unlabelled carbon in the medium is 1%, the fraction of labelled carbon will account to 99%. Hence, X = 0.99 | If the dilution by unlabelled carbon in the medium is 5%, the fraction of labelled carbon will account to 95%. Hence, X = 0.95 | If the dilution by unlabelled carbon in the medium is 10%, the fraction of labelled carbon will account to 90%. Hence, X = 0.9 |
|---|---|---|---|
| $M_1 = X (1-X)^{19} (^{20}C_1)$ | $1.98E^{-37}$ | $3.62E^{-24}$ | $1.8E^{-18}$ |
| $M_2 = X^2(1-X)^{18}(^{20}C_2)$ | $1.86E^{-34}$ | $6.54E^{-22}$ | $1.54E^{-16}$ |
| $M_3 = X^3 (1-X)^{17}(^{20}C_3)$ | $1.11E^{-31}$ | $7.46E^{-20}$ | $8.31E^{-15}$ |
| $M_4 = X^4 (1-X)^{16}(^{20}C_4)$ | $4.65E^{-29}$ | $6.02E^{-18}$ | $3.18E^{-13}$ |
| $M_5 = X^5 (1-X)^{15}(^{20}C_5)$ | $1.47E^{-26}$ | $3.66E^{-16}$ | $9.15E^{-12}$ |
| $M_6 = X^6 (1-X)^{14} (^{20}C_6)$ | $3.65E^{-24}$ | $1.74E^{-14}$ | $2.06E^{-10}$ |
| $M_7 = X^7 (1-X)^{13} (^{20}C_7)$ | $7.23E^{-22}$ | $6.61E^{-13}$ | $3.71E^{-09}$ |
| $M_8 = X^8 (1-X)^{12} (^{20}C_8)$ | $1.16E^{-19}$ | $2.04E^{-11}$ | $5.42E^{-08}$ |
| $M_9 = X^9 (1-X)^{11} (^{20}C_9)$ | $1.53E^{-17}$ | $5.17E^{-10}$ | $6.51E^{-07}$ |
| $M_{10} = X^{10}(1-X)^{10}(^{20}C_{10})$ | $1.67E^{-15}$ | $1.08E^{-08}$ | $6.44E^{-06}$ |
| $M_{11} = X^{11} (1-X)^9(^{20}C_{11})$ | $1.5E^{-13}$ | $1.87E^{-07}$ | $5.27E^{-05}$ |
| $M_{12} = X^{12} (1-X)^8(^{20}C_{12})$ | $1.12E^{-11}$ | $2.66E^{-06}$ | 0.0003 |
| $M_{13} = X^{13} (1-X)^7(^{20}C_{13})$ | $6.8E^{-10}$ | $3.11E^{-05}$ | 0.0019 |
| $M_{14} = X^{14} (1-X)^6(^{20}C_{14})$ | $3.37E^{-08}$ | 0.0002 | 0.0088 |
| $M_{15} = X^{15} (1-X)^5(^{20}C_{15})$ | $1.33E^{-06}$ | 0.0022 | 0.0319 |
| $M_{16} = X^{16} (1-X)^4(^{20}C_{16})$ | $4.13E^{-05}$ | 0.0133 | 0.0897 |
| $M_{17} = X^{17} (1-X)^3(^{20}C_{17})$ | 0.0009 | 0.0596 | 0.1901 |
| $M_{18} = X^{18} (1-X)^2(^{20}C_{18})$ | 0.0158 | 0.1886 | 0.2851 |
| $M_{19} = X^{19} (1-X) (^{20}C_{19})$ | 0.1652 | 0.3773 | 0.2701 |
| $M_{20} = X^{20}$ | 0.8179 | 0.3584 | 0.1215 |
| With the increase in dilution by unlabelled carbon source the corresponding X decreases and so does the probability of fully labelled camptothecin fraction. Moreover, as the dilution increases, the probability of camptothecin with a lower number of carbons getting labelled becomes higher. As shown above, at 10% dilution (X=0.9), the probability of 18 carbons getting labelled in the camptothecin ($M_{18}$) produced becomes highest (0.28), even more than $M_{20}$. This was also evident from the maximum relative abundance of $M_{18}$ observed during LC-MS analysis of the sample, obtained from D-[U-$^{13}$C]-glucose experiment. | | | |

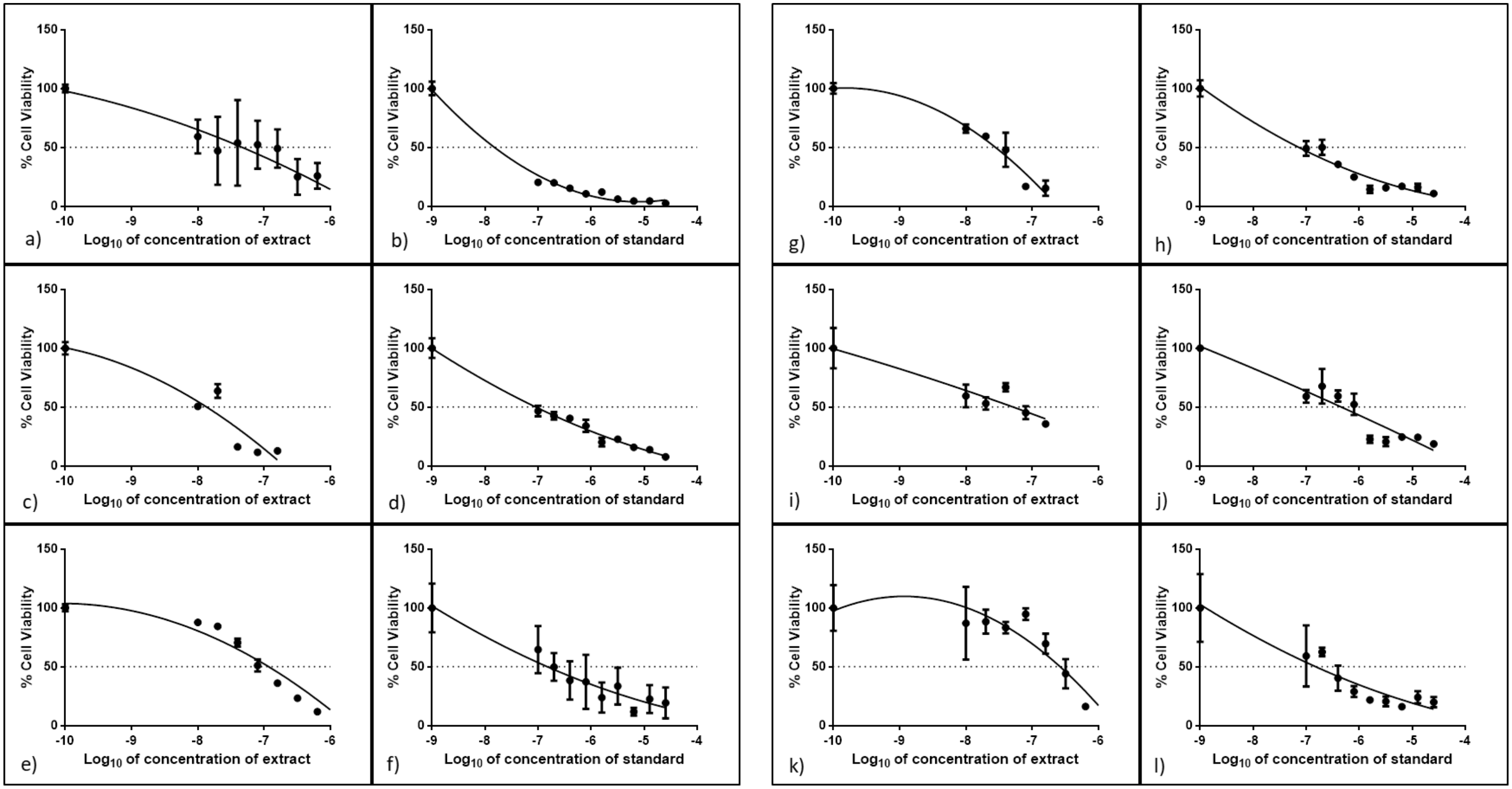


**Figure S6.** Effect of camptothecin (in extract and standard) on cell viability. Cytotoxic effects of camptothecin extract and the camptothecin standard at varying concentrations on MCF7 (a, b), H1299 (c, d), SKOV3 (e, f), Caco-2 (g, h), HT29 (i, j), and HEK293T (k, l) cell lines cell line

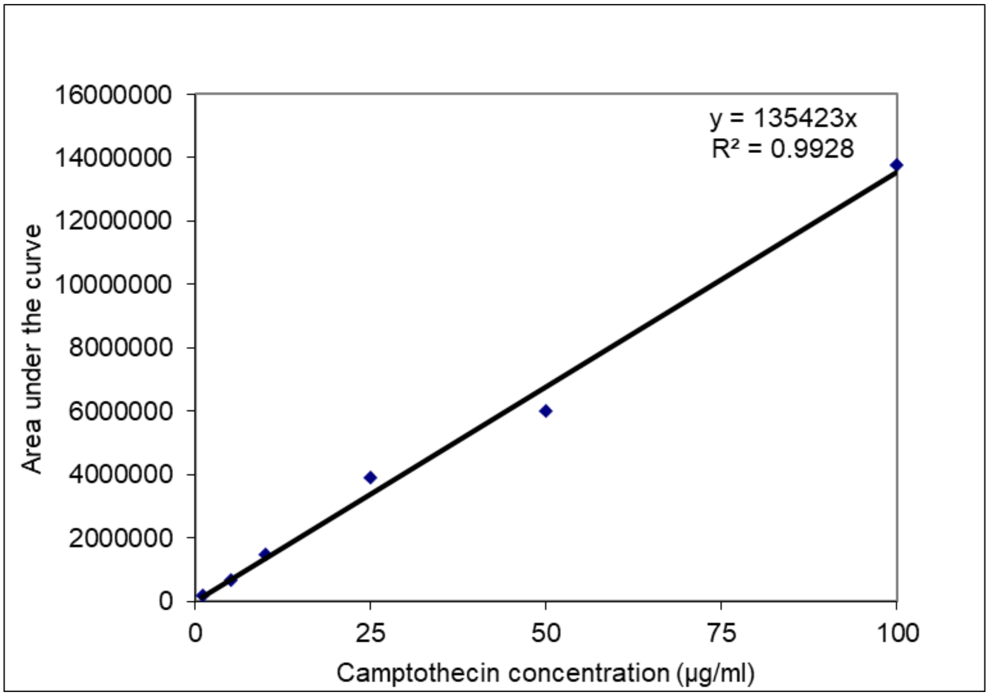


**Figure S7**. The standard curve of camptothecin obtained by plotting various concentrations of camptothecin against their respective peak area from HPLC.

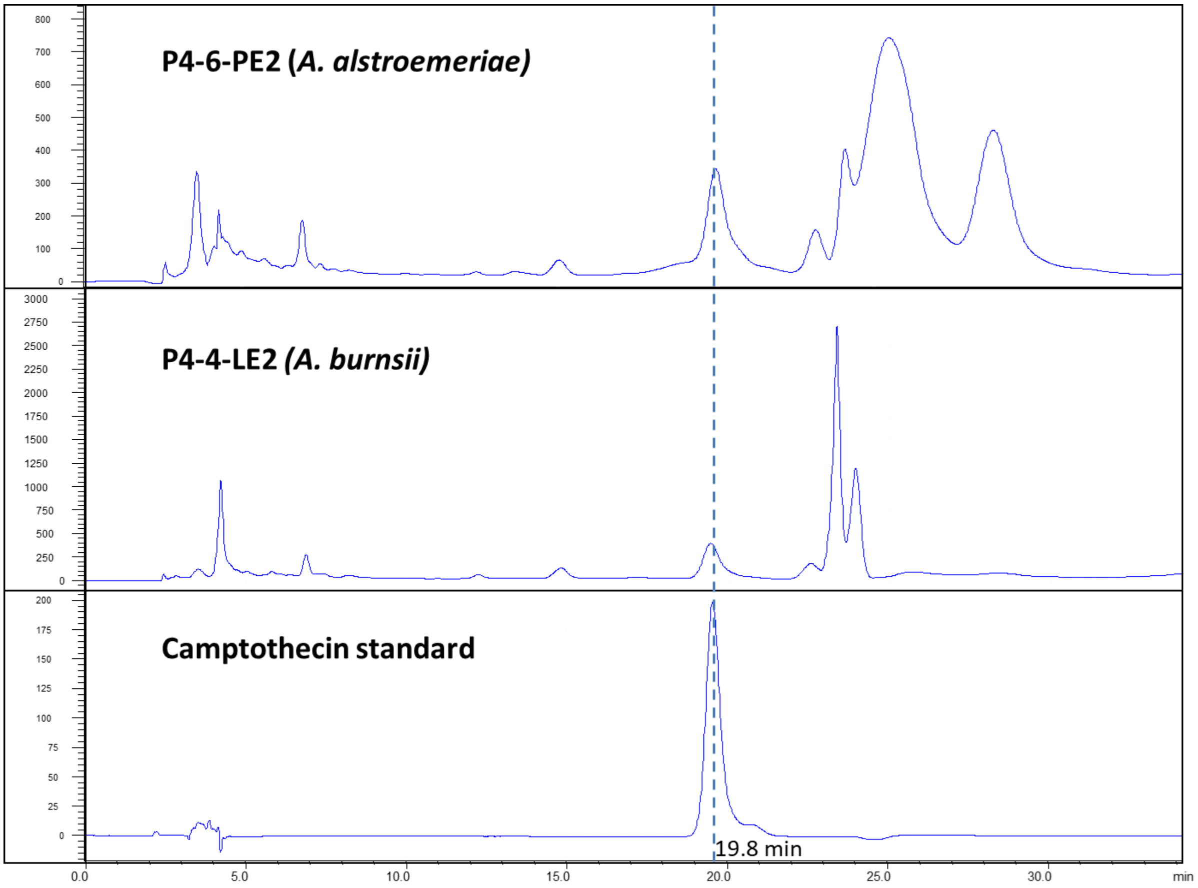


**Figure S8**. The chromatograms obtained from P4-6-PE2 (*A. alstroemeriae*) (top) and P4-4-LE2 (*A. burnsii*) (middle) using HPLC, compared with that of standard camptothecin (bottom).

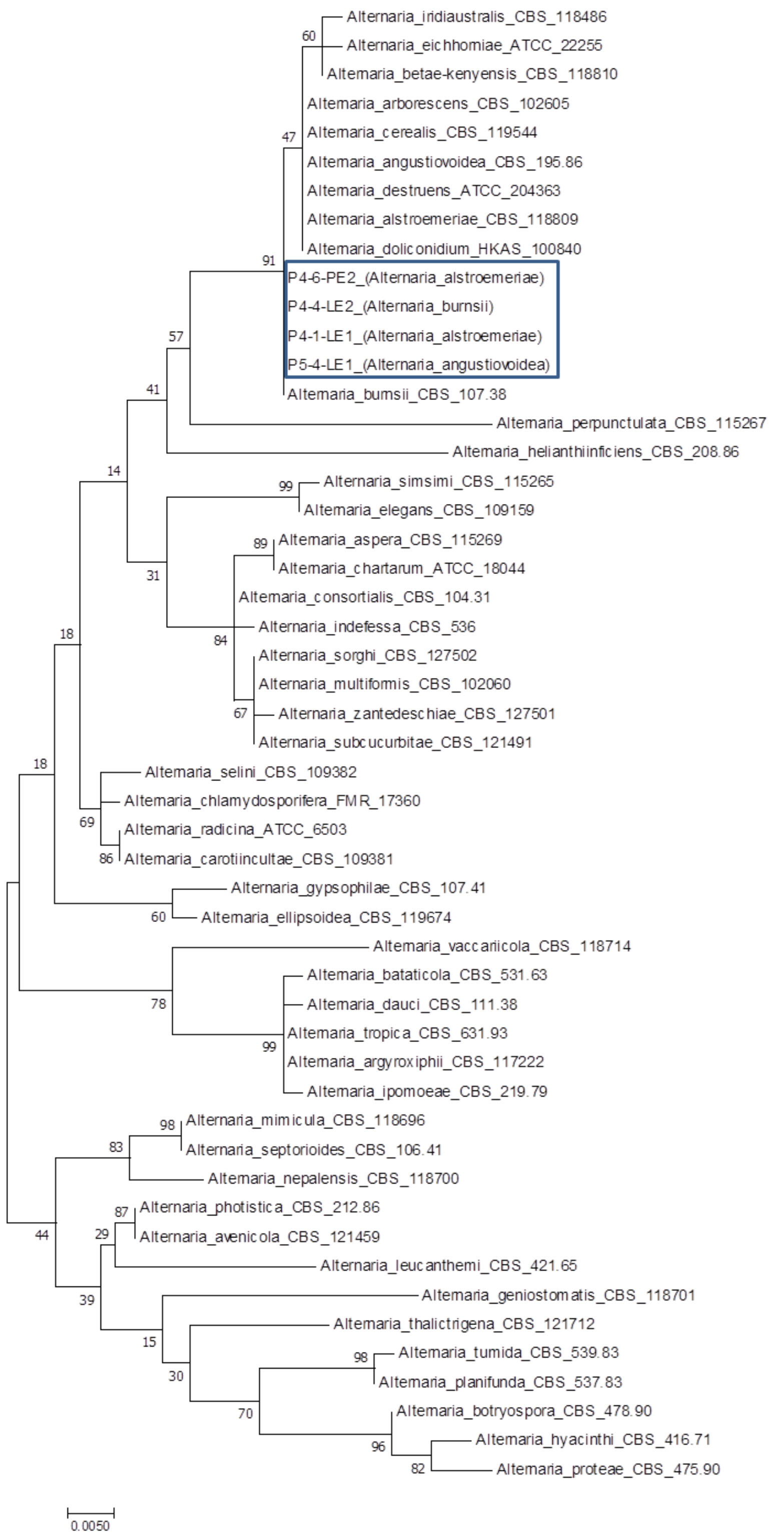


**Figure S9.** Molecular Phylogenetic analysis by Maximum Likelihood method based on the Jukes-Cantor model [28]. The strains isolated in this study are shown inside a blue rectangular box. The tree with the highest log likelihood (-1835.77) is shown. The percentage of trees in which the associated taxa clustered together is shown next to the branches. Initial tree(s) for the heuristic search were obtained automatically by applying Neighbor-Join and BioNJ algorithms to a matrix of pairwise distances estimated using the Maximum Composite Likelihood (MCL) approach, and then selecting the topology with superior log likelihood value. The tree is drawn to scale, with branch lengths measured in the number of substitutions per site. The analysis involved 51 nucleotide sequences. Codon positions included were 1st+2nd+3rd+Noncoding. All positions containing gaps and missing data were eliminated. There were a total of 471 positions in the final dataset. Evolutionary analyses were conducted in MEGA7. [29]
